# Adversarial Deconfounding Autoencoder for Learning Robust Gene Expression Embeddings

**DOI:** 10.1101/2020.04.28.065052

**Authors:** Ayse B. Dincer, Joseph D. Janizek, Su-In Lee

## Abstract

**Motivation:** Increasing number of gene expression profiles has enabled the use of complex models, such as deep unsupervised neural networks, to extract a latent space from these profiles. However, expression profiles, especially when collected in large numbers, inherently contain variations introduced by technical artifacts (e.g., batch effects) and uninteresting biological variables (e.g., age) in addition to the true signals of interest. These sources of variations, called confounders, produce embeddings that fail to transfer to different domains, i.e., an embedding learned from one dataset with a specific confounder distribution does not generalize to different distributions. To remedy this problem, we attempt to disentangle confounders from true signals to generate biologically informative embeddings.

**Results:** In this paper, we introduce the AD-AE (<u>A</u>dversarial <u>D</u>econfounding <u>A</u>uto<u>E</u>ncoder) approach to deconfounding gene expression latent spaces. The AD-AE model consists of two neural networks: (i) an autoencoder to generate an embedding that can reconstruct original measurements, and (ii) an adversary trained to predict the confounder from that embedding. We jointly train the networks to generate embeddings that can encode as much information as possible without encoding any confounding signal. By applying AD-AE to two distinct gene expression datasets, we show that our model can (1) generate embeddings that do not encode confounder information, (2) conserve the biological signals present in the original space, and (3) generalize successfully across different confounder domains. We demonstrate that AD-AE outperforms standard autoencoder and other deconfounding approaches.

**Availability:** Our code and data are available at https://gitlab.cs.washington.edu/abdincer/ad-ae.

## 1 Introduction

Gene expression profiles provide a snapshot of cellular activity, which allows researchers to examine the associations among expression, disease, and environmental factors. This rich information source has been explored by many studies, ranging from those that predict complex traits (Golub *et al.*, 1999; Shedden *et al.*, 2008; Geeleher *et al.*, 2014) to those that learn expression modules (Chun Tang *et al.*, 2001; Segal *et al.*, 2005; Teschendorff *et al.*, 2007). Advances in profiling technologies are rapidly increasing the availability of expression datasets. This has enabled the application of the complex non-linear models, such as neural networks, to various biological problems in order to identify signals not detectable using simple linear models (Lyu and Haque, 2018; Preuer *et al.*, 2018; Chaudhary *et al.*, 2018).

Unsupervised deep learning has enormous potential to extract important biological signals from the vast amount of expression profiles, as explored by recent studies (Tan *et al.*, 2016; Dincer *et al.*, 2018; Du *et al.*, 2019). Two features of unsupervised learning make it well suited to gene expression analysis. (i) *The ability to train informative models without supervision*, critical because it is challenging to obtain a high number of expression samples with coherent labels. Although many new expression profiles are released daily, the portion of the datasets with labels of interest is often too small. Moreover, different studies may collect information on different traits and even measure the same traits using different metrics (Haibe-Kains *et al.*, 2013). (ii) *The use of unsupervised models trained to extract patterns from the data without imposed directions or restrictions*. This aspect can be key to unlocking biological mechanisms yet unknown to the scientific community. Using unsupervised models to learn biologically meaningful representations would make it possible to map new samples to the learned space and adapt our model to any downstream task.

It is not straightforward to employ promising unsupervised models on gene expression data because expression measurements often contain out-of-interest sources of variation in addition to the signal we seek. When training an unsupervised model, we want the model to capture the true signal and learn latent dimensions corresponding to biological variables of interest. Especially when collected from a large cohort or multiple cohorts, expression profiles have, in addition to the true signal, variations in expression measures across samples as a result of (1) technical artifacts that are not relevant to biology, such as batch effects, (2) out-of-interest biological variables, such as sex, age, medications, and (3) random noise. (See Fig. 1.) We call these biological or non-biological artifacts that systematically affect expression values *confounders*. Unfortunately, in many datasets, confounder-based variations often mask true signals, which hinders learning biologically meaningful representations.

**Fig. 1.**
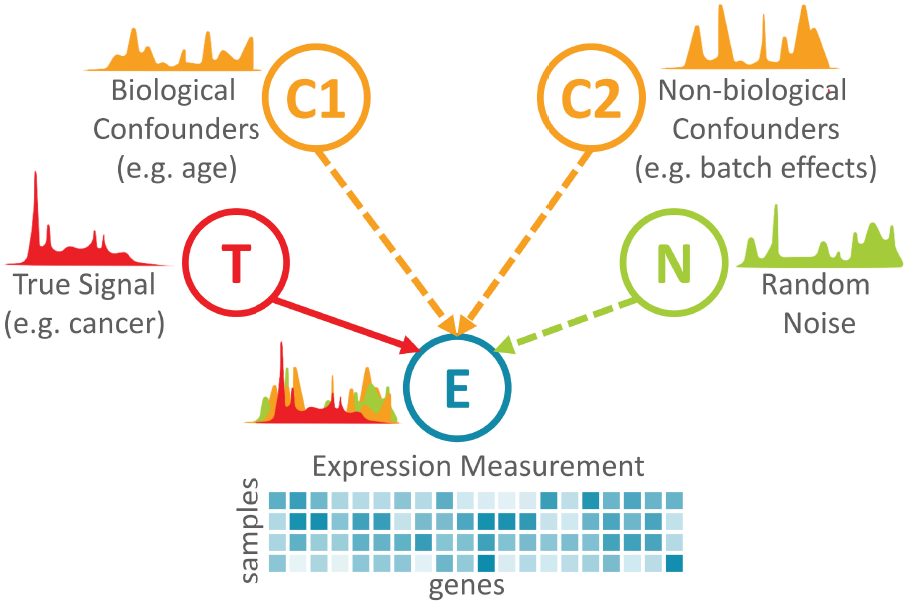
A simplified graphical model of measured expression shown as a mix of true signal, confounders of biological and non-biological origin, and random noise. Note that this model shows neither possible connections between a true signal and confounders nor connections among confounders.

As **a motivating example**, Fig. 2a shows how confounder signals might dominate true signals in gene expression data. We consider the KMPlot breast cancer expression dataset (Györffy *et al.*, 2010), which combines multiple microarray studies from The Gene Expression Omnibus (GEO) (Edgar *et al.*, 2002). We take the two GEO datasets with the highest number of samples and plot the first two principal components (PCs) (Wold *et al.*, 1987) to examine the strongest sources of variation. Fig. 2a shows that the two datasets are clearly separated, exemplifying how confounder-based variations affect expression measurements.

**Fig. 2.**
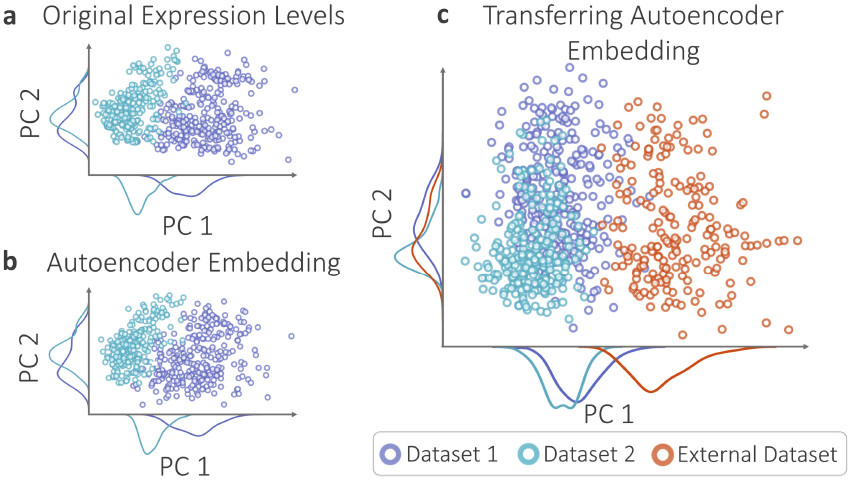
An example of confounder effects. The plot of top two principal components (PCs) colored by dataset labels generated for (a) the expression matrix, and (b) autoencoder embedding of the expression. (c) PC plot of the embeddings for training and external samples generated by the autoencoder trained from only the two datasets and transferred to the third external dataset.

**Fig. 3.**
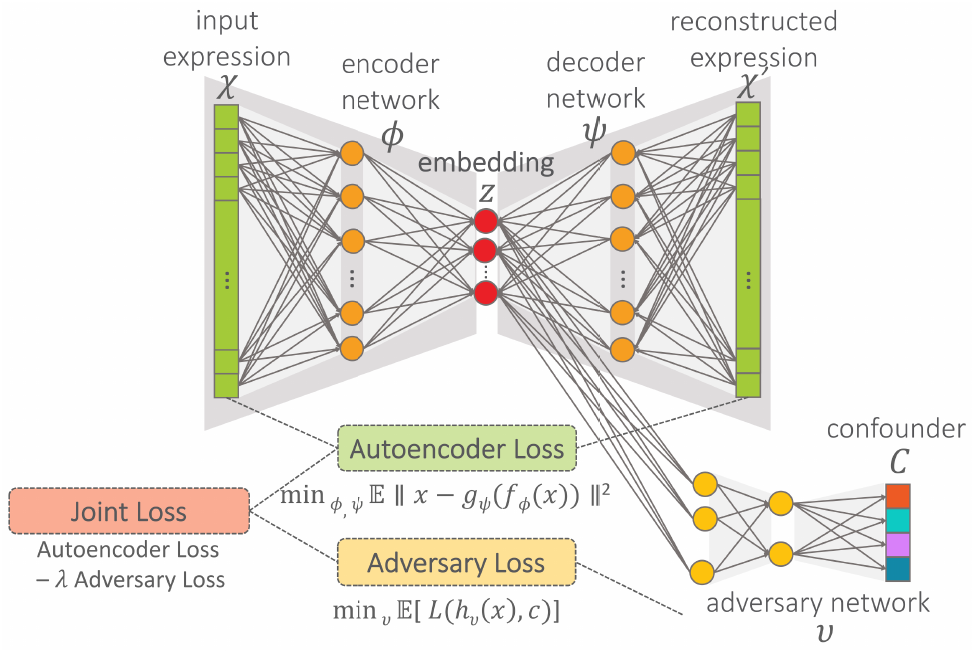
Adversarial deconfounding autoencoder (AD-AE) architecture. The model consists of an autoencoder and an adversary network. We jointly optimize the two models to minimize the joint loss, defined as the combination of reconstruction error and adversary loss.

We then apply an autoencoder (Hinton and Salakhutdinov, 2006) to this dataset, i.e., an unsupervised neural network that can learn a latent space that maps *M* genes to *D* nodes (*M* ≫ *D*) such that the biological signals present in the original expression space can be preserved in *D*-dimensional space. The autoencoder tries to capture the strongest sources of variation to reconstruct the original input successfully. In our example, unfortunately, it is encoding variation introduced by confounders rather than interesting signals. Fig. 2b depicts the PC plot of the autoencoder embedding. It shows that the dataset difference is encoded as the strongest source of variation. When we measure the Pearson’s correlation coefficient (Lin, 1989) between each node value and the binary dataset label, we observe that 78% of the embedding nodes are significantly correlated with the dataset label (p-value < 0.01). This means that most latent nodes are contaminated, making it difficult to disentangle biological signals from confounding ones.

Confounders also prevent our learning a robust, *transferable* model to generate generalizable embeddings that capture biological signals conserved across different domains. For instance, if we learn a model from one expression dataset that detects a disease signal, we want this signal to be valid for similar datasets. To simulate this problem, we use a separate set of samples from a different GEO study from the KMPlot data. We train the autoencoder using only the first two datasets, and we then encode the “external” samples from the third GEO study using the trained model. The PC plot in Fig. 2c highlights the distinct separation between the external dataset and the two training datasets. This simple example shows how confounder effects can prevent us from learning transferable latent models.

In this paper, we address the entanglement of confounders and true biological signals to show the power of deep unsupervised models to unlock biological mechanisms. Our goal is to generate biologically informative expression embeddings that are both robust to confounders and generalizable. To achieve this goal, we propose a deep learning approach to learning deconfounded expression embeddings, which we call AD-AE (<u>A</u>dversarial <u>D</u>econfounding <u>A</u>uto<u>E</u>ncoder).

AD-AE consists of two neural networks trained simultaneously: an autoencoder network optimized to generate an embedding that can reconstruct the data as successfully as possible, and (ii) an adversary network optimized to predict the confounder from the generated embedding. These two networks compete against each other to learn the optimal embedding that encodes important signals without encoding the variation introduced by the selected confounder variable. To demonstrate the performance of AD-AE, we used two expression datasets – breast cancer microarray and brain cancer RNA-Seq – with a variety of confounder variables, such as dataset label and age. We showed that AD-AE can generate unsupervised embeddings that preserve biological information while remaining invariant to selected confounder variables. We also conducted transfer experiments to demonstrate that AD-AE embeddings are generalizable across domains.

## 2 Methods

### 2.1 Standard Autoencoder

We used a standard autoencoder as the baseline for our experiments, which takes as input an expression vector *x* of *M* genes. The autoencoder consists of (i) an encoder network, defined as *f*_*ϕ*_ : *X* ↦ *Z*, which maps from the input space *X* ∈ ℝ^*M*^ to latent embedding *Z* ∈ ℝ^*D*^, and (ii) a decoder network, *g*_*ψ*_ : *Z* ↦ *X*, that maps the embedding *Z* back to the input space. Our encoder/decoder networks are fully or densely connected neural networks with rectified linear unit (ReLu) activation between layers; thus, *Z* is in effect the network’s information bottleneck. We then optimize over encoder and decoder networks as follows:

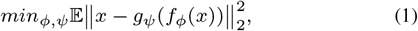

where *ϕ* and *ψ* are the parameters of our encoder and decoder neural networks, respectively. The expectation is taken over the training data, and the loss is the squared 2-norm distance between the input *x* and the reconstructed input.

### 2.2 Our Approach: Adversarial Deconfounding Autoencoder (AD-AE)

We propose the adversarial deconfounding autoencoder to generate biologically informative gene expression embeddings robust to confounders. AD-AE consists of two networks. The first is an autoencoder model *l* (defined in Section 2.1) that is optimized to generate an embedding that can reconstruct the original input. The second is an adversary model *h* that takes the embedding generated by the autoencoder as input and tries to predict the confounder *C*. We note that *C* is not limited to being a single confounder and could be a vector of them.

Our goal is to learn an embedding *Z* that encodes as much information as possible while not encoding any confounding signal. To achieve this, we train models *l* and *h* simultaneously. Model *l* tries to reconstruct the data while also *preventing the adversary from accurately predicting the confounder*. At the same time, adversarial predictor *h* tries to update its weights to accurately predict the confounder from the generated embedding. As shown by Louppe *et al.*, 2017, assuming the existence of an optimal model and sufficient statistical power, models *l* and *h* will converge and reach an equilibrium after a certain number of epochs, where *l* will generate an embedding *Z* that is optimally successful at reconstruction and *h* will only randomly predict a confounder variable from this embedding. In other words, the autoencoder will converge to generating an embedding that contains no information about the confounder, and the adversary will converge to a random prediction performance.

We train our model in three steps:

**Step 1:** The autoencoder model *l* is defined per Section 2.1. We pretrain the autoencoder to optimize equation 1 and generate an embedding *Z*.

**Step 2:** We define the adversary model *h*_*v*_ : *Z* ↦ *C*, mapping the embedding *Z* to confounder *C*. We again use fully connected multilayer perceptron networks with ReLu activation for the adversary, which is optimized with the following objective:

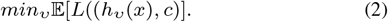

Here, we define a general loss function *L* that can be any differentiable function appropriate for the confounder variable (e.g., mean squared error for continuous confounders, cross-entropy for categorical confounders). We pretrain our adversary model accordingly to predict the confounder as successfully as possible.

**Step 3:** After separately pretraining both networks, we begin joint adversarial training by optimizing over the two networks. When optimizing the joint model, we first freeze the weights of the adversary model and train the autoencoder model for one epoch on a randomly selected minibatch of the data using stochastic gradient descent to optimize the following objective:

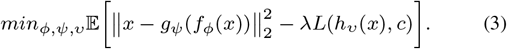

This corresponds to updating the weights of the autoencoder to *minimize* equation 1 while *maximizing* 2 (minimizing the negative of the objective). We then freeze the autoencoder model and train the adversary for an entire epoch to *minimize* equation 2. We continue this alternating training process until both models are optimized. If no model is simultaneously optimal at reconstructing the input expression without encoding confounding signals, the λ variable determines the ratio of weight the model gives to reconstruction or deconfounding. Increasing the λ value would learn a more deconfounded embedding while sacrificing reconstruction success; decreasing it would improve reconstruction at the expense of potential confounder involvement. For our experiments, we set λ = 1 since we believe this value maintains a reasonable balance between reconstruction and deconfounding.

## 3 Related Work

Though more general in scope, our paper is relevant to batch effect correction techniques. In high-throughput data, we often experience systematic variations in measurements caused by technical artifacts unrelated to biological variables, called *batch effects*. Many techniques have been developed to eliminate batch effects and *correct* high-throughput measurement matrices. Our work differs from batch correction approaches in two ways. First, we do not focus only on *batch effects*; rather we aim to build a model generalizable to any biological or non-biological confounder. Second, we do not concentrate on correcting the data, i.e., trying to eliminate confounder-sourced variations from the expression and outputting a *corrected* version of the expression matrix. Instead, our major objective is learning a confounder-free representation. We seek to reduce the dimension of an expression matrix in order to learn meaningful biological patterns that do not include confounders.

While keeping these differences in mind, we can compare our approach to batch correction techniques to highlight the advantages of our adversarial confounder-removal framework. In their review, Lazar *et al.*, 2012, categorize batch correction techniques into two groups. (i) Location-scale methods, which match the distribution of different batches by adjusting the mean and standard deviation of the genes. Examples include mean-centering (Sims *et al.*, 2008), gene-standardization (Li and Wong, 2001), ratio-based correction (Luo *et al.*, 2010), distance-weighted discrimination (Benito *et al.*, 2004), and probably the most popular of these techniques, the Empirical Bayes method (i.e., ComBat) (Johnson *et al.*, 2006). (ii) Matrix factorization techniques, which factorize the expression matrix to identify factors associated with batch effects and then reconstruct the data to eliminate batch-affected components. Examples include surrogate variable analysis (Leek and Storey, 2007) and various extensions of it (Teschendorff *et al.*, 2011; Parker *et al.*, 2014).

One limitation that applies to previously listed methods is that they model batch effects *linearly*. AD-AE, on the other hand, can eliminate *nonlinear* confounder effects as well. Several recent studies accounted for non-linear batch effects and tried modeling them with neural networks. These studies used either (i) maximum mean discrepancy (Borgwardt *et al.*, 2006) to match the distributions of two batches present in the data, such as Shaham *et al.*, 2017 and Amodio *et al.*, 2019, or (ii) an adversarial approach for batch removal, such as training an autoencoder with two separate decoder networks that correspond to two different batches along with an adversarial discriminator to differentiate the batches (Shaham, 2018) or generative adversarial networks trained to match distributions of samples from different batches (Upadhyay and Jain, 2019).

These methods all handle nonlinear batch effects. However, their application domain is limited since they can correct only for binary batch labels. AD-AE is a general model that can be used with any categorical or continuous valued confounder. To our knowledge, only Dayton, 2019 used an adversarial model to remove categorical batch effects, extending the approaches limited to binary labels. Our approach is significantly different since we focus on removing confounders from the latent space to learn deconfounded embeddings instead of trying to deconfound the reconstructed expression. Another unique aspect of our paper is that we concentrate on learning generalizable embeddings for which we carry transfer experiments for various expression domains and offer these domain transfer experiments as a new way of measuring the robustness of expression embeddings.

Our work takes its inspiration from research in *fair machine learning*, where the goal is to prevent models from unintentionally encoding information about sensitive variables, such as sex, race, or age. Multiple studies aimed to generate fair representations that try to learn as much as possible from the data without learning the membership of a sample to sensitive categories (Zemel *et al.*, 2013; Louizos *et al.*, 2015). Two studies with high relevance to our approach are Ganin *et al.*, 2016 and Louppe *et al.*, 2017, which use adversarial training to eliminate confounders. Ganin *et al.*, 2016 applied this idea to an autoencoder network to predict a class label of interest while avoiding encoding the confounder variable. Louppe *et al.*, 2017 also used an adversarial training approach by fitting an adversary model on the outcome of a classifier network to deconfound the predictor model. One advantage of Louppe’s model over the others is that it can work with any confounder variable, including continuous valued confounders. Janizek *et al.*, 2020 applied this approach to predict pneumonia from chest radiographs, showing that the model performs successfully without being confounded by selected variables.

Inspired by this work, we adopt a similar adversarial training approach for expression data, which is highly prone to confounders. Unlike prior work, AD-AE fits an adversary model on the *embedding space* to generate robust, confounder-free embeddings. In other words, we explore adversarial training to deconfound expression latent spaces in order to capture generalizable biological signals.

## 4 Experiments and Datasets

### 4.1 Datasets and Use Cases

We demonstrate the broad applicability of our model by employing it on two different expression datasets and experimenting with three different cases of confounders. Our method aims to both remove confounders from the embedding and encode as much biological signal as possible. Accordingly, we evaluate our model using two metrics: (i) *how successfully the embedding can predict the confounder*, where we expect a prediction performance close to random, and (ii) *the quality of prediction of biologically relevant variables*, where a better model is expected to lead to more accurate predictions.

Our first dataset was KMPlot (Györffy *et al.*, 2010), which offers a collection of breast cancer expression datasets from GEO microarray studies (Edgar *et al.*, 2002). We selected five GEO datasets with the highest number of samples from KMPlot, yielding a total of 1,139 samples and 13,018 genes (GEO accession numbers: GSE2034, GSE3494, GSE12276, GSE11121, and GSE7390). The confounder variable, the dataset label that was a categorical variable, indicated which of the five datasets each subset came from. For this dataset, we chose estrogen receptor (ER) and cancer grade as the biological variables of interest, since both are informative cancer traits. ER is a binary label that denotes the existence of estrogen receptors in cancer cells, an important phenotype for determining treatment (Knight *et al.*, 1977). Similarly, cancer grade can take values 1, 2, or 3 for invasive breast cancer, an indicator of the differentiation and growth speed of a tumor (Rakha *et al.*, 2010).

Our second dataset was brain cancer (glioma) RNA-Seq expression profiles obtained from TCGA, which contained lower grade glioma (LGG) and glioblastoma multiforme (GBM) samples (McLendon *et al.*, 2008; Brennan *et al.*, 2013; Brat *et al.*, 2015). We had a total of 672 samples and 20,502 genes. For this dataset, we used two different confounder variables as two separate use cases: sex as a binary confounder, and age as a continuous-valued one. For the biological trait, we used cancer subtype label, a binary variable indicating whether a patient had LGG or GBM, the latter a particularly aggressive subtype of glioma.

We preprocessed both datasets by applying standard gene expression preprocessing steps: mapping probe ids to gene names, log transforming the values, and making each gene zero-mean univariate. We also applied k-means++ clustering (Arthur and Vassilvitskii, 2006) on the expression data before training autoencoder models to reduce the number of features and decrease model complexity. We observed improvement in autoencoder performance when we applied clustering first and passed cluster centers to the model.

### 4.2 Deep Learning Architecture

For each individual dataset, we applied 5-fold cross validation to select the hyperparameters of autoencoder models. When training the model, we left out 20% of the samples for validation and determined the optimal number of epochs based on validation loss. We used the same autoencoder architecture for the AD-AE as well. Selection of the optimal number of latent nodes is beyond the scope of this paper; we tried to select a reasonable latent dimension size with respect to the number of samples we had. Yet to demonstrate that our model is invariant to the latent dimension size, we experimented with various sizes ranging from 10 to 100 nodes.

For the breast cancer data, we extracted 1,000 k-means cluster centers since the input dimension was slightly above 1,000. The latent space size was set to 100, resulting in a 10% dimension reduction. Our selected model had one hidden layer in both encoder and decoder networks, with 500 hidden nodes and a dropout rate of 0.1. The minibatch size was 128, and we trained with Adam optimizer (Kingma and Ba, 2014) using a learning rate of 0.0005. ReLU activation was applied to all layers of the encoder and decoder except the last layer, where we applied linear activation. For the adversarial model, we used a fully connected neural network that had 2 hidden layers with 100 hidden nodes in each layer, and we used ReLU activation. The last layer had 5 hidden nodes corresponding to the number of confounder classes and softmax activation. The adversarial model was trained with categorical cross entropy loss.

The architecture selected for brain cancer expression was very similar, with 500 k-means cluster centers, 50 latent nodes, one hidden layer with 500 nodes in both networks with no dropout, and ReLU activation at all layers except the last layers of the networks; the remaining parameters were the same as those for the breast cancer network. The adversarial model was also the same except for 50 hidden nodes in each layer. For the sex confounder, the last layer had 1 hidden node with sigmoid activation, trained with binary cross entropy loss; for the age confounder, the last layer used linear activation, trained with mean squared loss.

We implemented AD-AE using Keras with Tensorflow background.

### 4.3 Alternative Approaches to Deconfounding

When evaluating our model, the most straightforward competitor was a standard autoencoder, which allowed us to directly observe the effects of confounder removal. We also compared against other commonly used approaches to confounder removal. For all these different techniques, we first applied the correction method and then trained an autoencoder model to generate an embedding from the corrected data. We could not compare against nonlinear batch effect correction techniques (Section 3) since they were applicable only on binary confounder variables.

#### Batch mean-centering

(Sims *et al.*, 2008) subtracts the average expression of all samples from the same confounder class (e.g., batch) from the expression measurements.

#### Gene standardization

(Li and Wong, 2001) transforms each gene measurement to have zero mean and one standard deviation within a confounder class.

#### Empirical Bayes method (ComBat)

(Johnson *et al.*, 2006) matches distributions of different batches by mean and deviation adjustment. To estimate the mean and standard deviation for each confounder class, the model adopts a parametric or a non-parametric approach to gather information about confounder effects from groups of genes with similar expression patterns.

## 5 Results

### 5.1 Adversarial Deconfounding Autoencoder Learns Biologically Meaningful Deconfounded Embeddings

Our first experiment aimed to demonstrate that AD-AE could successfully encode the biological signals we wanted while not detecting the selected confounders. We used the KMPlot breast cancer expression dataset and trained standard autoencoder and AD-AE to create embeddings, and we generated PC plots showing the two orthogonal dimensions with the highest variation (Fig. 4). Observe that the standard autoencoder embedding clearly separates datasets, indicating that the learned embedding was highly confounded (Fig. 4a i). On the other hand, the PC plot for AD-AE embedding shows that data points from different datasets are fused (Fig. 4b i). This is expected since we trained our model until both networks converged, which means that we obtained a random prediction performance on the validation set for the adversarial network. More interestingly, we colored the PC plots by biological variables of interest: ER status and cancer grade. Observe that for the autoencoder embedding, the samples are not differentiated by phenotype labels (Fig. 4a ii&iii). This shows that when we learn an embedding with a standard autoencoder model, confounders might dominate the embedding, preventing it from learning clear biological patterns. On the other hand, the PC plot of the AD-AE embedding clearly distinguishes samples by ER label as well as cancer grade (Fig. 4b ii & iii), showing the effects of deconfounding. We further investigate these results in Section 5.3 by fitting prediction models on the embeddings to quantitatively evaluate the models.

**Fig. 4.**
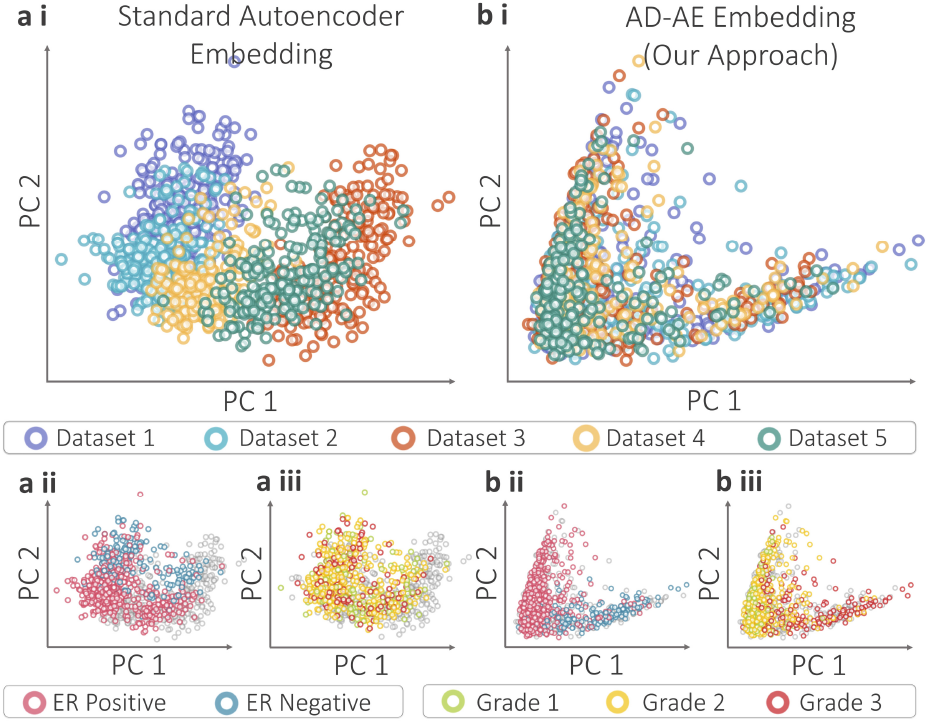
Principal Component (PC) plots of embeddings generated by (a) standard autoencoder, and (b) Adversarial Deconfounding Autoencoder (AD-AE). Subplots are colored by (i) dataset, (ii) ER status, and (iii) cancer grade. The gray dots denote samples with missing labels.

We created the same plots for the standard AE and AD-AE embeddings, this time using UMAP (McInnes *et al.*, 2018), which is replacing t-SNE (Maaten and Hinton, 2008) for generating interpretable visualizations of data (Fig. 5). We demonstrate that the UMAP plot sharply distinguishes the datasets from the autoencoder embedding, while AD-AE does not separate the samples by dataset label. This again lets us identify ER labels and cancer grade from the AD-AE embedding, though it is not possible to reasonably differentiate selected biological variables from the standard autoencoder embedding.

**Fig. 5.**
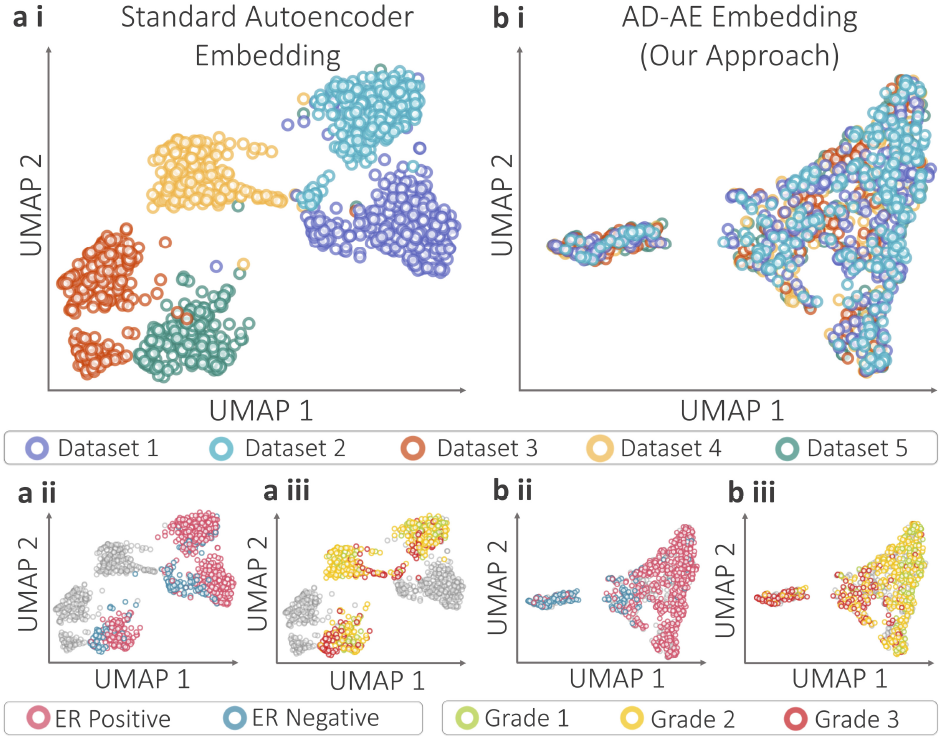
UMAP plots of embeddings generated by (a) standard autoencoder, and (b) Adversarial Deconfounding Autoencoder (AD-AE). Subplots are colored by (i) dataset, (i) ER status, and (iii) cancer grade. The gray dots denote samples with missing labels.

### 5.2 AD-AE Can Learn Embeddings Generalizable to Different Domains

AD-AE generates embeddings that are robust to confounders and generalizable to different domains. The most common applications for this model are learning an embedding from a dataset and transferring it to a separate dataset. To simulate this problem with breast cancer samples, we left one dataset out for testing and trained the standard autoencoder on the remaining four datasets. We then generated two embeddings for the internal and external datasets: (i) one for samples from the four datasets used for training, and (ii) another for the left out samples from the fifth dataset. Note that we trained the model using samples in the four datasets only, and we then used the already trained model to encode the fifth dataset samples. We then repeated the same training and encoding procedure for AD-AE to compare the generalizability of both models.

In Fig. 6, the circle and diamond markers denote the UMAP representation of the embedding generated for training and left-out dataset samples, respectively. In Fig. 6a i, we colored all samples by their ER labels. First of all, we draw attention to the external set data points that are clustered entirely separately from the training samples. This shows that the standard embedding does not precisely generalize to left-out samples. More importantly, we do not a see a general direction of separation for the ER labels that is valid for both the training and left-out samples; (ER+ samples are clustered on the right for training samples and mainly on the left for external samples). This clustering indicates that the manifold learned for the training samples does not transfer to the external dataset. We observed the same scenario when we colored the same plots by cancer grade (Fig. 6a ii).

**Fig. 6.**
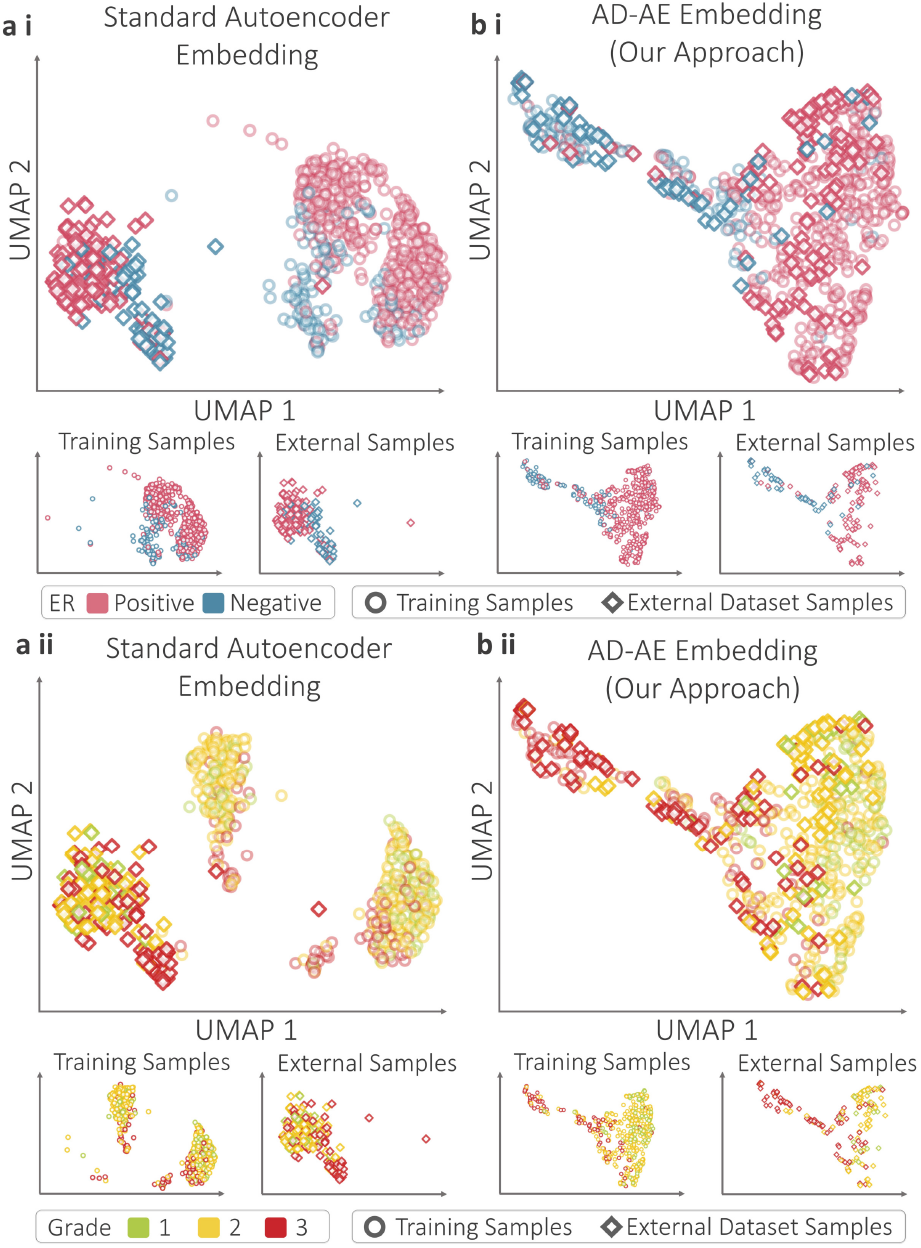
UMAP plots of embeddings generated by (a) standard autoencoder, and (b) Adversarial Deconfounding Autoencoder. Plots are colored by (i) Estrogen Receptor (ER) labels, and (ii) cancer grade labels. The circle and diamond markers denote training and external dataset samples, respectively. For clarity, the subplots for the training and external samples are provided below the joined plots.

To examine whether the adversarial deconfounding autoencoder can better generalize to a separate dataset, we created UMAP plots (as in Fig. 6a) for the AD-AE embedding (Fig. 6b). We emphasize that it is not possible to distinguish training from external samples because the circle and diamond markers overlap one another. But the critical point is the separation of samples by ER label (Fig. 6b i). Observe that ER-samples from the training set are concentrated on the upper left of the plot, while ER+ samples dominate the right. The same direction of separation applies to the samples from the external dataset. This plot concisely demonstrates that when we remove confounders from the embedding, we can learn generalizable biological patterns otherwise overshadowed by confounder effects. We applied the same analysis using the cancer grade labels and again observed the same pattern (Fig. 6b ii).

### 5.3 AD-AE Can Successfully Predict Biological Phenotypes

To show that AD-AE preserves the true biological signals present in the expression data, we predicted cancer phenotypes from the learned embeddings. In Sections 5.1 and 5.2, we visualized our embeddings to demonstrate how our approach removes confounder effects and learns meaningful biological representations. Nonetheless, we wanted to offer a quantitative analysis as well to thoroughly compare our model to a standard baseline and to alternative deconfounding approaches. After generating embeddings with AD-AE and competitor models, we fit prediction models to the embeddings to predict biological phenotypes of interest. We also applied the prediction test on different domains to examine how well the learned embeddings generalized to external test sets and measure the generalization gap for each model as a metric of robustness.

In Fig. 7a, we show the ER prediction performance of our model compared to all other baselines. To predict ER status, we used an elastic net classifier, tuning the regularization and l1 ratio parameters with 5-fold cross validation. We recorded the area under precision-recall curves (PR-AUC) since the labels were unbalanced. We separately selected the optimal model for each embedding generated by AD-AE and each competitor. To measure each method’s consistency, we repeated the embedding generation process 10 times with *10 independent random trainings of the models*, and we ran prediction tasks for each of the 10 embeddings for each model. We used linear models for the prediction for two reasons. First, the sample size was small due to the missingness of phenotype labels for some samples and the splitting of samples across domains, which made it difficult to fit complex models. Second, reducing the expression matrix dimension size let us reduce complexity and fit simpler models to capture patterns.

**Fig. 7.**
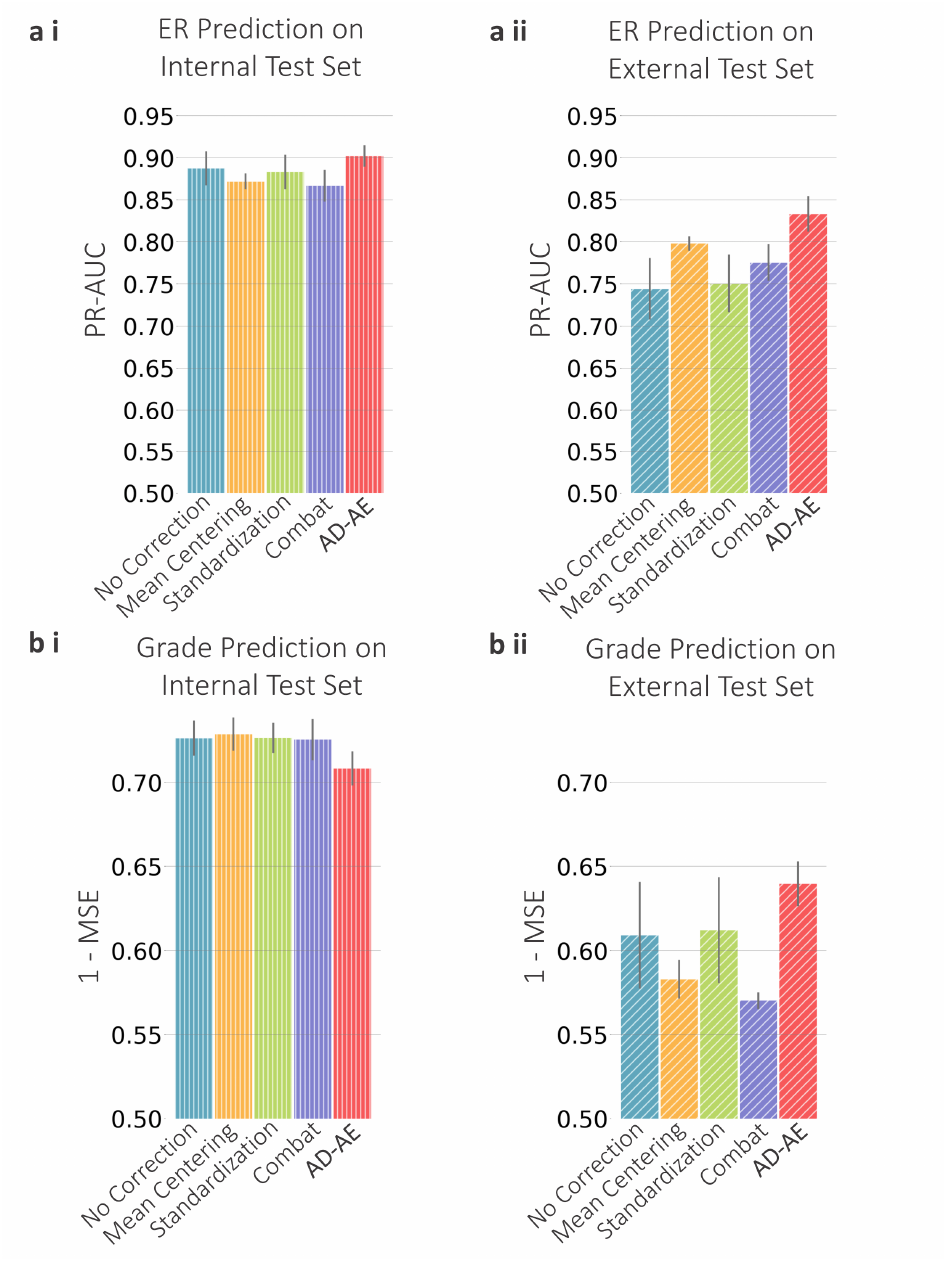
(a) Estrogen Receptor (ER) prediction plots for (i) internal test set, and (ii) external test set. (b) Cancer grade prediction plots.

We trained AD-AE and the competitors using only four datasets, leaving the fifth dataset out. To train the linear prediction model, we left out 20% of the samples from the four datasets for testing, trained the model using the rest of the samples, then predicted on the left-out *internal* samples to measure PR-AUC. To measure prediction performance of the external dataset, we used the exact same training samples obtained from the four datasets and then predicted for the *external* dataset samples. In Fig. 7a i, observe that for the internal dataset, our model barely outperforms other baselines and the uncorrected model. This is expected: when the domain is the same, we might not see the advantage of confounder removal. However, Fig. 7a ii shows that when predicting for the left-out dataset, AD-AE clearly outperforms all other models. This result shows that AD-AE much more successfully generalizes to other domains.

We repeated the same experiments, this time to predict cancer grade, for which we fit an elastic net regressor tuned with 5-fold cross validation, measuring the mean squared error. Fig. 7b shows that for the internal prediction, our model is not as successful as other models; however, it outperforms all baselines in terms of external test set performance. This result indicates that a modest decrease in internal test set performance could significantly improve our model’s external test set performance.

Moreover, we showed that the generalization gap of AD-AE is much smaller than the baselines we compare against (Fig. 8). We calculated the generalization gap as the distance between internal and external test set prediction scores. A high generalization gap means that model performance declines sharply when transferred to another domain; a small generalization gap indicates a model can transfer across domains with minimal performance decline. Therefore, AD-AE successfully learns manifolds that are valid across different domains, as we demonstrated for both ER and cancer grade predictions.

**Fig. 8.**
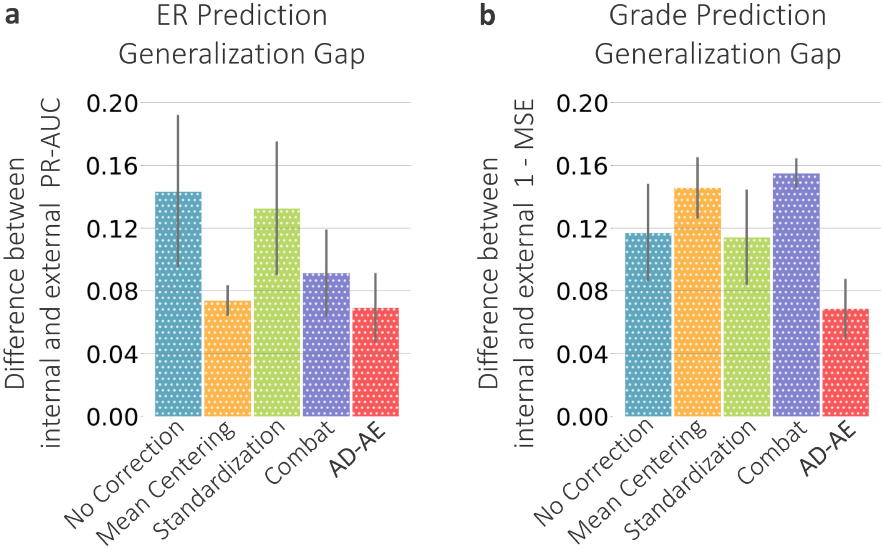
The generalization gap calculated as the difference between the internal and external prediction scores shown for (a) ER prediction and (b) cancer grade prediction.

### 5.4 AD-AE Embeddings Can Be Successfully Transferred Across Domains

We next extend our experiments to the TCGA brain cancer dataset to further evaluate AD-AE. We trained our model and the baselines with the same procedure we applied to the breast cancer dataset and again fitted prediction models. We first trained an elastic net classifier to predict cancer subtype (LGG vs GBM) from the embeddings. We trained the predictor model using only female samples and predicted for male samples. We then repeated this transfer process, this time training from male samples and predicting on females. Note that the autoencoder was trained from all samples (male and female), and prediction models were trained from one class of samples (e.g., males) and transferred to another class (e.g., females). This experiment was intended to evaluate how accurate an embedding would be at predicting biological variables of interest when the confounder domain is changed. Fig. 9 shows that AD-AE easily outperforms the standard baseline and all competitors for both transfer directions.

**Fig. 9.**
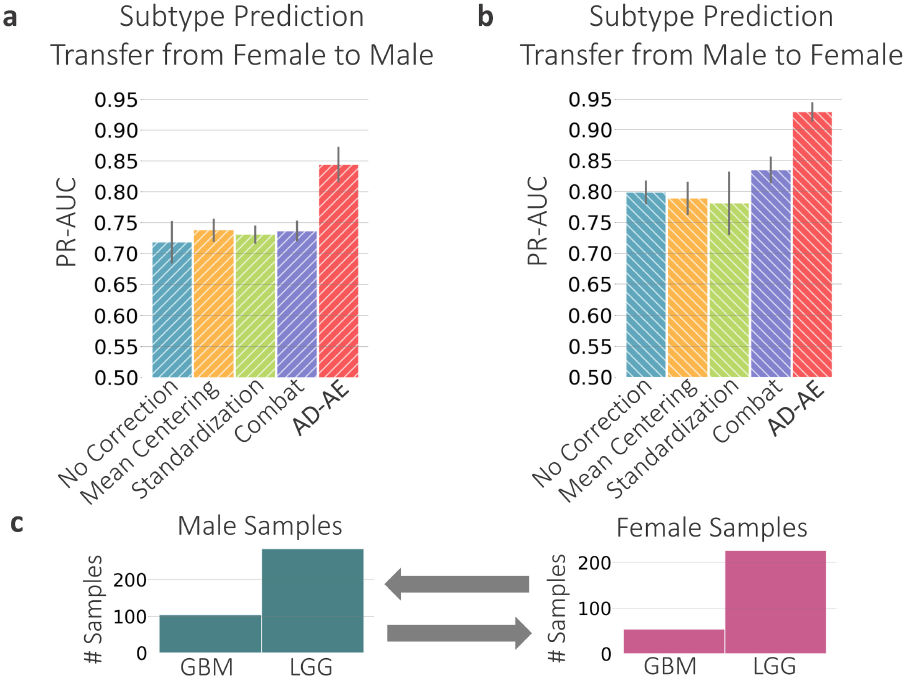
Glioma subtype prediction plots for (a) model trained on female samples transferred to male samples and (b) model trained on male samples transferred to female samples. (c) Subtype label distributions for male and female samples.

We find this result extremely promising since we offer confounder domain transfer prediction as a metric for evaluating the robustness of an expression embedding. Researchers want to generate informative embeddings that encode biological signals without being confounded by variables (e.g., sex). We note that the confounder variable is data and domain dependent, and sex can be a crucial biological variable of interest for certain diseases or datasets. In this experiment, we wanted to learn about cancer subtypes and severity independent of a patient’s sex. We succeed at this task of accurately predicting complex phenotypes regardless of the distribution of the confounder variable. We also highlight that our model can solve the problem of class imbalance that commonly occurs in domain shift (Ming Harry Hsu *et al.*, 2015). Fig. 9c shows that the distribution of cancer subtypes differs for male and female domains. This might lead to discrepancies when transferring from one domain to another; however, AD-AE embeddings could be successfully transferred independent of the distribution of labels, a highly desirable property of a robust expression embedding.

We repeated the same experiments using age as the continuous-valued confounder variable. Other models were not applicable for continuous valued confounders; thus, we can compare only to the standard baseline. For the prediction transfer experiments, we again fit an elastic net classifier to predict cancer subtype and separated the samples into two groups: samples with age within one standard deviation (i.e., center of the distribution), and samples with age beyond one standard deviation (i.e., edges of the distribution). Fig. 10c shows the age distribution of the brain cancer dataset, highlighting the samples in the center and on the edges. We trained the predictor on the center samples and predicted for samples on the edge, and vice versa (Fig. 10a & b). Our model substantially outperforms the standard baseline in both transfer directions. Especially, when we trained on samples within one standard deviation and predicted for remaining samples, we can see a huge increase in performance compared to the standard baseline. This case simulates a substantial age distribution shift. It is promising to see that disentangling confounders from expression embeddings can be the key to capturing signals generalizable over different domains, such as different age distributions.

**Fig. 10.**
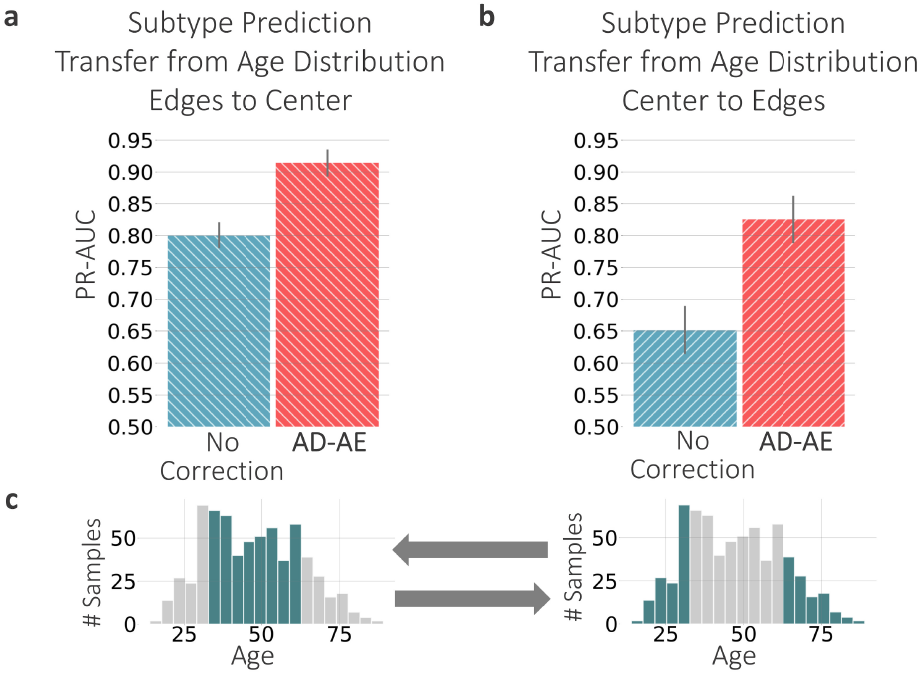
Glioma subtype prediction plots for (a) model trained on samples beyond one standard deviation of the age distribution (i.e., edges of the distribution) transferred to samples within one standard deviation (i.e., center of the distribution), and (b) vice versa. (c) Age distributions of all samples.

## 6 Discussion

Gene expression datasets contain valuable information central to unlocking biological mechanisms and understanding the biology of complex diseases. Unsupervised learning aims to encode information present in vast amounts of unlabeled samples to an informative latent space, helping researches discover signals without biasing the learning process.

Hindering the learning of meaningful representations is the fact that gene expression measurements often contain unwanted sources of variation, such as experimental artifacts and out-of-interest biological variables. Variations introduced by *confounders* can overshadow the true expression signal, preventing the model from learning accurate patterns. Particularly when we combine multiple expression datasets to increase statistical power, we can learn an embedding that encodes dataset differences rather than biological signals shared across multiple datasets.

We introduced the Adversarial Deconfounding Autoencoder (AD-AE) to generate expression embeddings robust to confounders. AD-AE trains two neural networks simultaneously, an autoencoder to generate an embedding that reconstructs the original data successfully and an adversary model that predicts the selected confounders from the generated embedding. We jointly optimized the two models; the autoencoder tries to learn an embedding free from the confounder variable, while the adversary tries to predict the confounder accurately. On convergence, the encoder learns a latent space where the confounder cannot be predicted even using the optimally trained adversary network.

We evaluated our model based on (1) deconfounding of the learned latent space, (2) preservation of biological signals, and (3) prediction of biological variables of interest when the embedding is transferred from one confounder domain to another. We experimented with two datasets, KMPlot breast cancer expression, where we used dataset labels as the confounder variable, and TCGA brain cancer RNA-Seq expression, where we used both sex and age as separate confounders. For these different use cases, we showed that AD-AE generates deconfounded embeddings that successfully predict biological phenotypes of interest. Importantly, we showed the advantage of our model over standard autoencoder and alternative deconfounding approaches on transfer experiments, where our model generalized much better to different domains.

A potential limitation of our approach is that we extend an unregularized autoencoder model by incorporating an adversarial component. We can improve our model by adopting a regularized autoencoder such as denoising autoencoder (Vincent *et al.*, 2008), or variational autoencoder (Kingma and Welling, 2013). In this way, we could prevent model overfitting and make our approach more applicable to datasets with smaller sample sizes. Another limitation is that although our model can train an adversary model to predict a vector of confounders, we have not yet conducted experiments to correct for multiple confounders simultaneously. We could extend our model by incorporating multiple adversarial networks to account for various confounders.

AD-AE is an adversarial approach for generating confounder-free embeddings for gene expression that can be easily adapted for any confounder variable. In this paper, we tested our model on cancer expression datasets since cancer expression samples are available in large numbers. However, we would like to extend testing to other expression datasets as well, including samples from different diseases and normal tissues. We also see as future work experimenting on single cell RNA-Seq data to learn informative embeddings combining multiple datasets.

## Acknowledgements

This work was supported by the National Institutes of Health [R35 GM 128638, and R01 NIA AG 061132]; National Science Foundation [CAREER DBI-1552309, and DBI-1759487]; and American Cancer Society [127332-RSG-15-097-01-TBG]. We are grateful to Sandy Kaplan for providing feedback on the manuscript. We also thankfully acknowledge all members of the AIMS lab for their helpful comments and useful discussions.

## References

Amodio, M. et al. (2019). Exploring single-cell data with deep multitasking neural networks. Nature Methods, 16(11), 1139–1145.

Arthur, D. and Vassilvitskii, S. (2006). k-means++: The advantages of careful seeding. Technical Report 2006-13, Stanford InfoLab.

Benito, M. et al. (2004). Adjustment of systematic microarray data biases. Bioinformatics, 20(1), 105–114.

Borgwardt, K. M. et al. (2006). Integrating structured biological data by Kernel Maximum Mean Discrepancy. Bioinformatics, 22(14), e49–e57.

Brat, D. J. et al. (2015). Comprehensive, integrative genomic analysis of diffuse lower-grade gliomas. New England Journal of Medicine, 372(26), 2481–2498.

Brennan, C. W. et al. (2013). The somatic genomic landscape of glioblastoma. Cell, 155(2), 462–477.

Chaudhary, K., Poirion, O. B., Lu, L., and Garmire, L. X. (2018). Deep learning–based multi-omics integration robustly predicts survival in liver cancer. Clinical Cancer Research, 24(6), 1248–1259.

Chun Tang, Li Zhang, Aidong Zhang, and Ramanathan, M. (2001). Interrelated two-way clustering: an unsupervised approach for gene expression data analysis. In Proceedings 2nd Annual IEEE International Symposium on Bioinformatics and Bioengineering (BIBE 2001), pages 41–48.

Dayton, J. B. (2019). Adversarial Deep Neural Networks Effectively Remove Nonlinear Batch Effects from Gene-Expression Data. Master’s thesis, Brigham Young University.

Dincer, A. B., Celik, S., Hiranuma, N., and Lee, S.-I. (2018). Deepprofile: Deep learning of cancer molecular profiles for precision medicine. bioRxiv.

Du, J. et al. (2019). Gene2vec: Distributed representation of genes based on co-expression. BMC Genomics, 20(82).

Edgar, R., Domrachev, M., and Lash, A. E. (2002). Gene Expression Omnibus: NCBI gene expression and hybridization array data repository. Nucleic Acids Research, 30(1), 207–210.

Ganin, Y. et al. (2016). Domain-adversarial training of neural networks. The Journal of Machine Learning Research, 17(59), 35.

Geeleher, P., Cox, N. J., and Huang, R. S. (2014). Clinical drug response can be predicted using baseline gene expression levels and in vitro drug sensitivity in cell lines. Genome Biology, 15(3), R47.

Golub, T. R. et al. (1999). Molecular classification of cancer: Class discovery and class prediction by gene expression monitoring. Science, 286(5439), 531–537.

Györffy, B. et al. (2010). An online survival analysis tool to rapidly assess the effect of 22,277 genes on breast cancer prognosis using microarray data of 1,809 patients. Breast Cancer Research and Treatment, 123(3), 725–731.

Haibe-Kains, B. et al. (2013). Inconsistency in large pharmacogenomic studies. Nature, 504(7480), 389–393.

Hinton, G. E. and Salakhutdinov, R. R. (2006). Reducing the Dimensionality of Data with Neural Networks. Science, 313(5786), 504–507.

Janizek, J. D., Erion, G., DeGrave, A. J., and Lee, S.-I. (2020). An adversarial approach for the robust classification of pneumonia from chest radiographs. arXiv preprint arXiv:2001.04051.

Johnson, W. E., Li, C., and Rabinovic, A. (2006). Adjusting batch effects in microarray expression data using empirical Bayes methods. Biostatistics, 8(1), 118–127.

Kingma, D. P. and Ba, J. (2014). Adam: A method for stochastic optimization. arXiv preprint arXiv:1412.6980.

Kingma, D. P. and Welling, M. (2013). Auto-encoding variational bayes. arXiv preprint arXiv:1312.6114.

Knight, W. A., Livingston, R. B., Gregory, E. J., and McGuire, W. L. (1977). Estrogen receptor as an independent prognostic factor for early recurrence in breast cancer. Cancer Research, 37(12), 4669–4671.

Lazar, C. et al. (2012). Batch effect removal methods for microarray gene expression data integration: a survey. Briefings in Bioinformatics, 14(4), 469–490.

Leek, J. T. and Storey, J. D. (2007). Capturing heterogeneity in gene expression studies by surrogate variable analysis. PLoS Genetics, 3(9), 1724–1735.

Li, C. and Wong, W. H. (2001). Model-based analysis of oligonucleotide arrays: Expression index computation and outlier detection. Proceedings of the National Academy of Sciences, 98(1), 31–36.

Lin, L. I.-K. (1989). A concordance correlation coefficient to evaluate reproducibility. Biometrics, 45(1), 255–268.

Louizos, C., Swersky, K., Li, Y., Welling, M., and Zemel, R. (2015). The variational fair autoencoder. arXiv preprint arXiv:1511.00830.

Louppe, G., Kagan, M., and Cranmer, K. (2017). Learning to pivot with adversarial networks. In Advances in Neural Information Processing Systems 30, pages 981–990.

Luo, J. et al. (2010). A comparison of batch effect removal methods for enhancement of prediction performance using MAQC-II microarray gene expression data. The Pharmacogenomics Journal, 10(4), 278–291.

Lyu, B. and Haque, A. (2018). Deep learning based tumor type classification using gene expression data. In Proceedings of the 2018 ACM International Conference on Bioinformatics, Computational Biology, and Health Informatics, pages 89–96.

Maaten, L. v. d. and Hinton, G. (2008). Visualizing data using t-sne. Journal of Machine Learning Research, 9(Nov), 2579–2605.

McInnes, L., Healy, J., and Melville, J. (2018). Umap: Uniform manifold approximation and projection for dimension reduction. arXiv preprint arXiv:1802.03426.

McLendon, R. et al. (2008). Comprehensive genomic characterization defines human glioblastoma genes and core pathways. Nature, 455(7216), 1061–1068.

Ming Harry Hsu, T. et al. (2015). Unsupervised domain adaptation with imbalanced cross-domain data. In The IEEE International Conference on Computer Vision (ICCV), pages 4121–4129.

Parker, H. S. et al. (2014). Preserving biological heterogeneity with a permuted surrogate variable analysis for genomics batch correction. Bioinformatics, 30(19), 2757–2763.

Preuer, K. et al. (2018). DeepSynergy: predicting anti-cancer drug synergy with Deep Learning. Bioinformatics, 34(9), 1538–1546.

Rakha, E. A. et al. (2010). Breast cancer prognostic classification in the molecular era: the role of histological grade. Breast Cancer Research, 12(4), 207.

Segal, E., Pe’er, D., Regev, A., Koller, D., and Friedman, N. (2005). Learning module networks. Journal of Machine Learning Research, 6(Apr), 557–588.

Shaham, U. (2018). Batch effect removal via batch-free encoding. bioRxiv.

Shaham, U. et al. (2017). Removal of batch effects using distribution-matching residual networks. Bioinformatics, 33(16), 2539–2546.

Shedden, K. et al. (2008). Gene expression–based survival prediction in lung adenocarcinoma: a multi-site, blinded validation study. Nature Medicine, 14(8), 822–827.

Sims, A. H. et al. (2008). The removal of multiplicative, systematic bias allows integration of breast cancer gene expression datasets–improving meta-analysis and prediction of prognosis. BMC Medical Genomics, 1(42).

Tan, J., Hammond, J. H., Hogan, D. A., and Greene, C. S. (2016). Adage-based integration of publicly available pseudomonas aeruginosa gene expression data with denoising autoencoders illuminates microbe-host interactions. mSystems, 1(1).

Teschendorff, A. E., Miremadi, A., Pinder, S. E., Ellis, I. O., and Caldas, C. (2007). An immune response gene expression module identifies a good prognosis subtype in estrogen receptor negative breast cancer. Genome Biology, 8(8), R157.

Teschendorff, A. E., Zhuang, J., and Widschwendter, M. (2011). Independent surrogate variable analysis to deconvolve confounding factors in large-scale microarray profiling studies. Bioinformatics, 27(11), 1496–1505.

Upadhyay, U. and Jain, A. (2019). Removal of batch effects using generative adversarial networks. arXiv preprint arXiv:1901.06654.

Vincent, P., Larochelle, H., Bengio, Y., and Manzagol, P.-A. (2008). Extracting and composing robust features with denoising autoencoders. In Proceedings of the 25th International Conference on Machine Learning, page 1096–1103.

Wold, S., Esbensen, K., and Geladi, P. (1987). Principal component analysis. Chemometrics and Intelligent Laboratory Systems, 2(1-3), 37–52.

Zemel, R., Wu, Y., Swersky, K., Pitassi, T., and Dwork, C. (2013). Learning fair representations. In International Conference on Machine Learning, pages 325–333.

